# Circadian distribution of epileptiform discharges in epilepsy: candidate mechanisms of variability

**DOI:** 10.1101/2022.08.24.505077

**Authors:** Isabella Marinelli, Jamie J. Walker, Udaya Seneviratne, Wendyl D’Souza, Mark J. Cook, Clare Anderson, Andrew P. Bagshaw, Stafford L. Lightman, Wessel Woldman, John R. Terry

## Abstract

Epilepsy is a serious neurological disorder characterised by a tendency to have recurrent, spontaneous, seizures. Classically, seizures are assumed to occur at random. However, recent research has uncovered underlying rhythms both in seizures and in key signatures of epilepsy - so-called interictal epileptiform activity - with timescales that vary from hours and days through to months. Understanding the physiological mechanisms that determine these rhythmic patterns of epileptiform discharges remains an open question. Many people with epilepsy identify precipitants of their seizures, the most common of which include stress, sleep deprivation and fatigue. To quantify the impact of these physiological factors, we analysed 24-hour EEG recordings from a cohort of 107 people with idiopathic generalized epilepsy. We found two subgroups with distinct distributions of epileptiform discharges: one with highest incidence during sleep and the other during day-time. We interrogated these data using a mathematical model that describes the transitions between background and epileptiform activity in large-scale brain networks. This model was extended to include a time-dependent forcing term, where the excitability of nodes within the network could be modulated by other factors. We calibrated this forcing term using independently-collected human cortisol (the primary stress-responsive hormone characterised by circadian and ultradian patterns of secretion) data and sleep-staged EEG from healthy human participants. We found that either the dynamics of cortisol or sleep stage transition, or a combination of both, could explain most of the observed distributions of epileptiform discharges. Our findings provide conceptual evidence for the existence of underlying physiological drivers of rhythms of epileptiform discharges. These findings should motivate future research to explore these mechanisms in carefully designed experiments using animal models or people with epilepsy.

**Author summary:** 65 million people have epilepsy worldwide. Many of these people report specific triggers that make their seizures (the primary symptom of epilepsy) more likely. Here, we use a mathematical model to understand the relationship between possible triggers and rhythms in epileptiform activity observed across the day.

The mathematical model describes the activity of connected brain regions, and how the excitability of these regions can change in response to different stimuli. Based on data collected from people with idiopathic generalized epilepsy, we identify transitions between sleep stages and variation in concentration of the stress-hormone cortisol as candidate factors that influence how likely it is for epileptiform activity to occur. By including those factors into the model, we show they can explain most of the daily variability. More broadly, our approach provides a framework for better understanding what factors drive the occurrence of epileptiform activity and offers the potential to suggest experiments that can validate model predictions.

## Introduction

Epilepsy is a common neurological disorder, affecting 65 million people globally [1–3]. The primary symptom of epilepsy – seizures – is believed to occur as a result of disruptions in the level of neuronal excitability. In particular, mechanisms that govern the normal balance between excitation and inhibition can become compromised causing parts of the brain to become hyperexcitable, which can be characterised at different scales. For example, at the cellular level it is strongly associated with the so-called paroxysmal depolarization shift (PDS) of cortical pyramidal cells [4, 5]. At the macroscale, it manifests in pathological electrical activity, captured using electroencephalography (EEG), called epileptiform discharges (EDs). EDs can be thought of as an umbrella term that encompasses both interictal (i.e., between seizures) epileptiform activity (e.g., spikes) as well as ictal activity (i.e., seizures).

Epileptiform activity has classically been thought to occur at random, but recent studies have presented compelling evidence for underlying rhythmicity in EDs [6–9]. Although such cycles have been shown to follow several temporal scales, including ultradian, circadian, multidien and even circannual rhythms [10, 11], relatively little is currently known about the mechanisms governing these rhythms and how intrinsic and extrinsic factors can modulate the likelihood of EDs. This limits the extent to which this knowledge of rhythmicity can be used for clinical benefit.

Many people with epilepsy identify triggers that appear to make them more likely, and some of these triggers are physiological factors known to influence cortical excitability. The most common of these are stress, sleep, hormones, and medication [12–17].

In this study, we consider some of these factors as candidate mechanisms that modulate the likelihood of EDs, and therefore provide insight into the mechanisms underlying observed distributions of EDs [17].

The mammalian stress-response is driven by circulating glucocorticoid hormones: predominantly cortisol in humans and corticosterone in rodents, herein CORT. Ultradian and circadian rhythms of CORT are controlled by the hypothalamic-pituitary-adrenal (HPA) axis, a neuroendocrine axis, wherein a delayed negative-feedback loop mediates hormone secretion from the pituitary and adrenal glands [18]. The impact of CORT on brain function is well established. For example, rapid changes in CORT secretion not only have major effects on glucocorticoid receptor activation in the brain [19] but also major effects on cognition [20]. Furthermore, Karst et al. [21] demonstrated that neuronal excitability is rapidly and reversibly determined by changes in CORT levels. At the macroscale, Schridde et al. [22] observed a CORT dose-dependent increase in EDs in the genetically in-bred Wistar Albino Glaxo/Rij (WAG-Rij) model of human idiopathic generalized epilepsy (IGE). A similar relationship has been found more recently in people with stress-sensitive focal epilepsies [23].

One of the most direct ways of measuring human cortical excitability is via transcranial magnetic stimulation (TMS), with motor and/or EEG responses taken as a proxy for excitability. With this approach, prolonged wakefulness leading to sleep deprivation has been shown to increase excitability or alter the excitatory-inhibitory balance of the supplementary motor cortex [24–26]. In addition, TMS-derived cortical excitability is also modulated by circadian phase, such that excitability is lowest in the early evening prior to bedtime, and peaks at the end of the biological night [27]. These observations have not always been consistent [28] with some suggestion of differences between participants with and without epilepsy [28]. These results are generally consistent with the changing probability of EDs associated with sleep deprivation and/or fluctuations in the circadian rhythm [29, 30]. The probability of EDs also varies through the sleep cycle, with non-rapid eye movement (NREM) sleep generally having a facilitatory effect, and REM sleep an inhibitory effect [30–32]. The latter observation is consistent with the increase in TMS-defined excitability associated with selective REM sleep deprivation [33].

However, the complexity of these interrelating factors, alongside the difficulty of simultaneously measuring their physiological correlates, makes unpacking them challenging. In this paper, we analysed distributions of EDs from 107 people with IGE collected over 24-hours. We found evidence to support the existence of two primary groups with different mechanisms driving the overnight likelihood of EDs and their likelihood during the day. To explore possible contributing factors underpinning these different mechanisms we developed a mathematical modelling framework that:

a. describes transitions between background states and EDs;
b. relates excitability to the likelihood of these transitions;
c. considers the impact of intrinsic and extrinsic factors on excitability.

We calibrated model parameters using independently collected 24-hour hormone profiles from 6 healthy participants, and sleep staged polysomnography data from 42 healthy participants. We used synthetic minority oversampling to account for discrepancies in group size, enabling us to generate synthetic distributions of EDs. We explored the goodness of fit between these model derived distributions and those observed in the cohort of people with IGE. Our mathematical analysis revealed evidence to support the view that the likelihood of EDs is modulated by both transitions in sleep stages, as well as by ultradian fluctuations in cycling CORT levels.

## Results

We analysed distributions of EDs derived from 24-hour EEG recordings from 107 subjects with IGE (see Materials and Methods for a detailed description of this data-set).

### Variability in the circadian distribution of epileptiform discharges

We found that the median number of EDs over 24 hours was approximately 29, although several individuals had more than 200 events (Fig 1, Panel A and B). Examination of normalised ED patterns on an hourly basis (i.e., for each individual, the number of EDs at each hour was divided by their total number of EDs and we then normalised over the cohort), suggested that the likelihood of EDs varied across the day.

**Fig 1.**
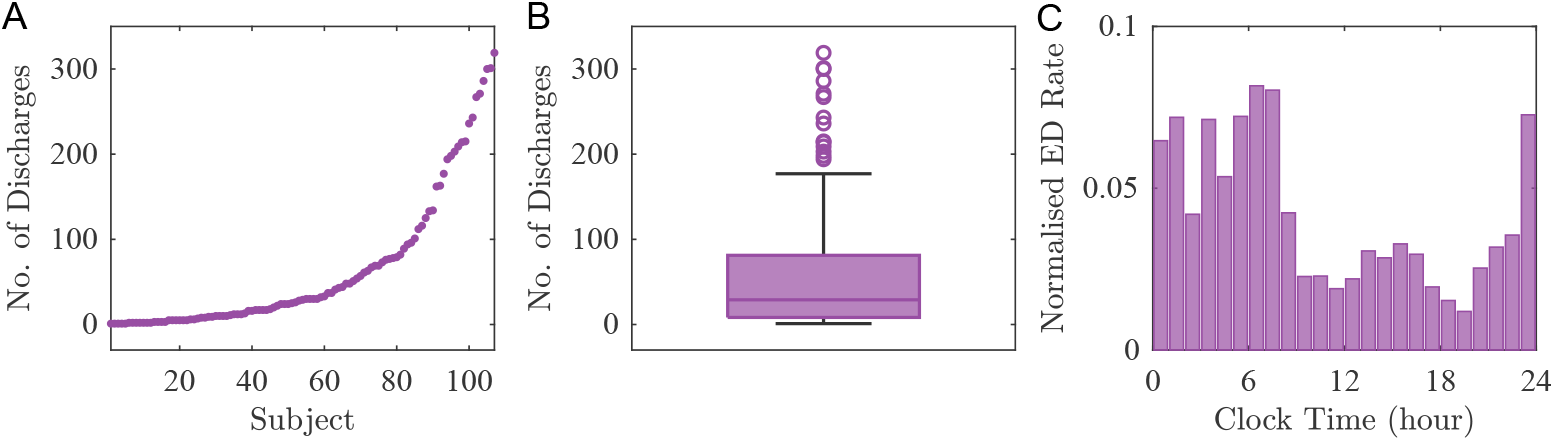
ED distribution in people with IGE. (A) Number of EDs from 107 subjects with idiopathic generalized epilepsy (IGE). (B) Boxplot shows basic sample statistics (minimum, lower quartile, median, upper quartile and maximum) of the number of EDs. (C) normalised EDs rate per hour.

To investigate the possible temporal distribution of EDs across the 24-hour day (herein referred to as the ‘circadian distribution’), we first considered similarities between subjects. We used MATLAB R2021a (MathWorks Inc., Natick, MA) to compute the cross-correlation coefficients of time series representing the individual hourly ED rate. This leads to a correlation matrix *C*, with entries *C*_*ij*_ corresponding to the similarity between the pattern of EDs in subject *i* and in subject *j* (Fig 2, A). The closer the value of *C*_*ij*_ to 1, the more similar the distribution of EDs of subject *i* and subject *j*. Subsequently, we clustered subjects according to their correlation coefficients using k-means clustering [34] and the Calinski-Harabasz criterion [35] to optimise the number of clusters (see S1 Appendix). This analysis revealed two primary groups within the overall cohort of people with IGE that displayed different temporal ED distribution patterns: Group 1 of 66 individuals and Group 2 of 41 individuals (Fig 2, B). Importantly, the identified clusters were found to be consistent across a range of bin widths (45-90 minutes) (see S2 Appendix).

**Fig 2.**
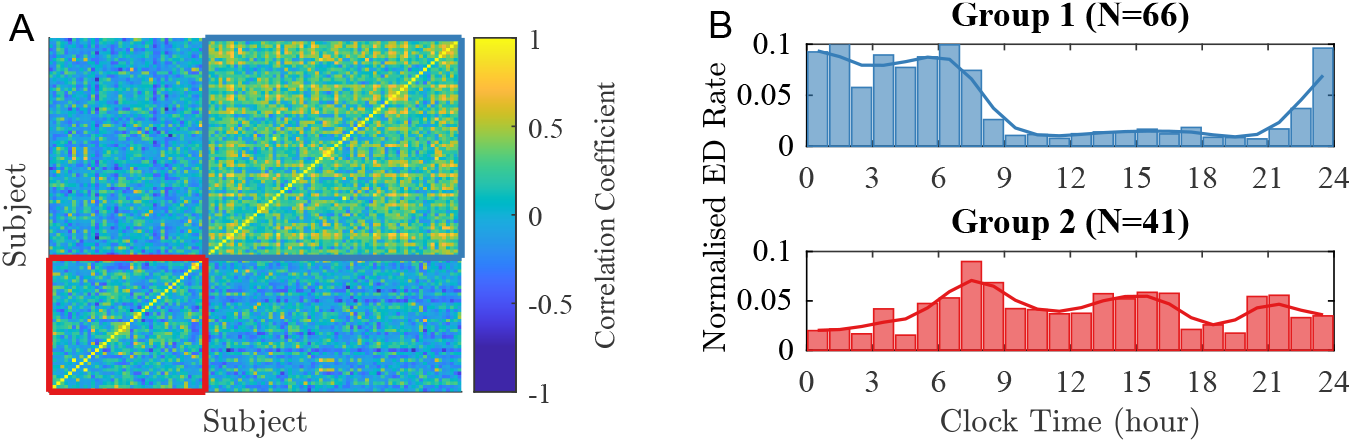
IGE subjects organised based on different circadian ED distribution patterns. (A) A pairwise cross-correlation matrix (of size 107 × 107) was calculated using ED hourly rate patterns in order to establish similarities within the IGE cohort. (B) Group 1 (blue, N = 66) and Group 2 (red, N = 41) were identified based on the similarities of hourly ED rate.

We found that the groups identified by our cluster analysis were not caused by imbalances in the type of epilepsy. Specifically, individuals with IGE were classified into childhood absence epilepsy (CAE), juvenile absence epilepsy (JAE), juvenile myoclonic epilepsy (JME), generalized epilepsy with generalized tonic-clonic seizures only (GTCSO), and genetic generalized epilepsy unspecified (GGEU) according to the criteria published by the ILAE [36]. We fitted a linear model (using R version 4.0.2) to assess the dependence of the groups on epilepsy type (see S3 Appendix for details), finding no evidence of an association (*p* = 0.193: two-tailed *t*-test).

### Candidate mechanisms impacting the distributions: sleep and CORT

We explored candidate mechanisms that could explain differences in ED distributions between the two groups identified by our cluster analysis. The (empirical) likelihood of EDs in Group 1 (Fig 2, B top) displayed a significant increase in the propensity for EDs during the night and lower levels during day-time. In contrast, the likelihood of EDs in Group 2 (Fig 2, B bottom) displayed greater variation during waking hours.

To assess the impact of inter-individual timing of sleep and its duration on ED distributions, we adjusted time within each subject such that *t* = 0 corresponded to either their sleep onset or sleep offset. The resulting distributions are presented in Fig 3 Panels A-D. For Group 1, we found that the ED rate was higher for approximately 9 hours starting at habitual sleep onset (Panel A), while it was relatively low during the rest of the day. In Panel B, we observed the same trend but shifted to the 9 hours before waking. For Group 2 (Fig 3, C and D) we did not find increased levels of EDs during sleep; instead, the distribution suggests a potential daytime ultradian rhythm.

**Fig 3.**
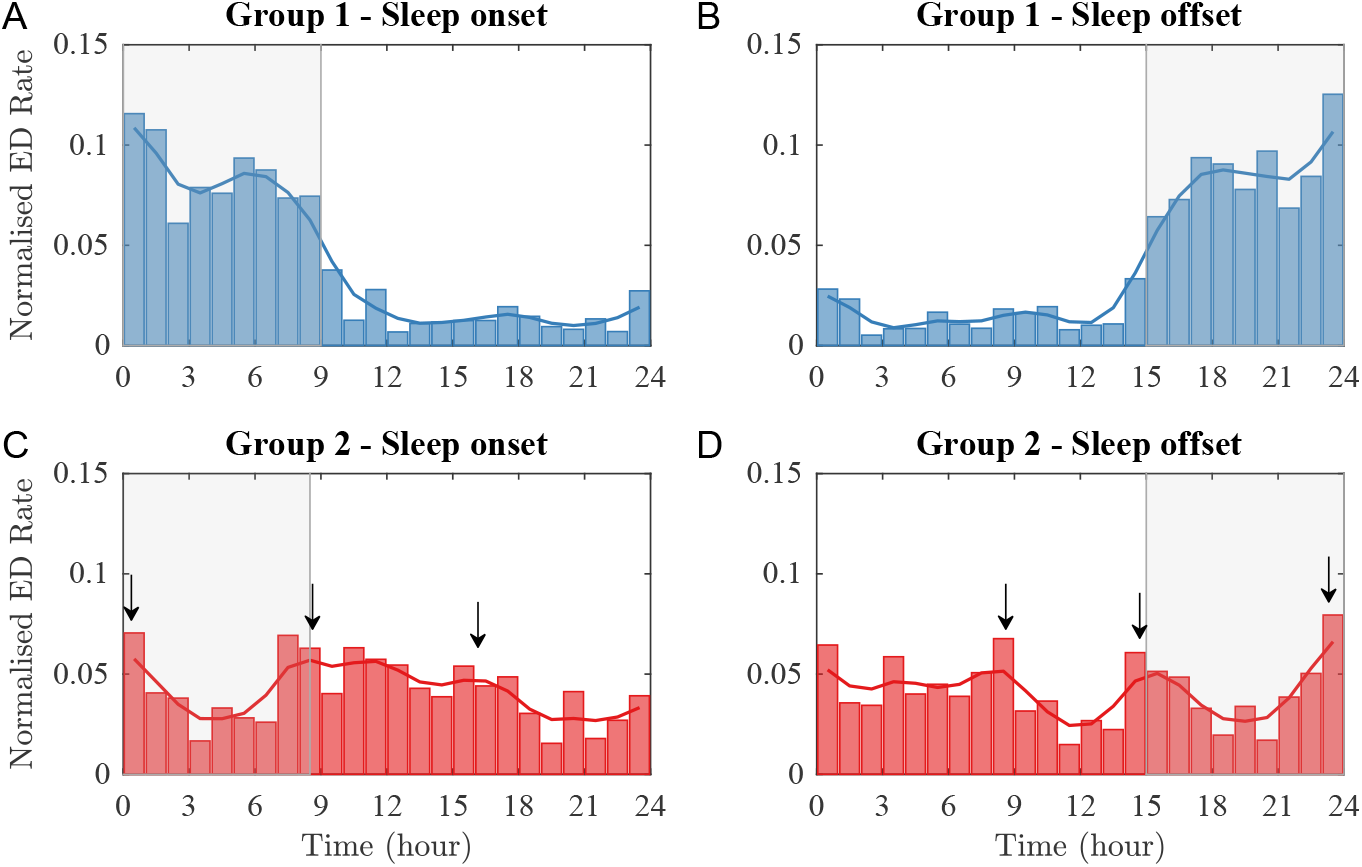
Impact of timing of sleep and its duration on ED distributions. Epileptiform discharges for Group 1 (top row) and Group 2 (bottom row) with time normalised such that *t* = 0 corresponds with sleep onset (A and C) and with sleep offset (B and D). The transparent grey box highlights the average habitual sleep period.

To quantify this more explicitly, we introduced the parameter *F*_*i*_, *i* = 1, 2 to measure the fraction of EDs occurring during sleep for each group:

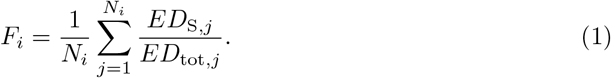

Here *N*_*i*_ is the number of subjects in Group *i, ED*_S,*j*_ and *ED*_tot,*j*_ are the numbers of ED occurrences for the *j*^*th*^ subject in the *i*^*th*^ group occurring during during the individual’s sleep time and across the full 24-hour period, respectively. We found *F*_1_ = 0.8, suggesting that 80% of EDs in Group 1 were clustered during the sleep period. In contrast *F*_2_ is 0.37, suggesting that in Group 2 just over a third of discharges occur during the sleep period, consistent with the 8-9 hour sleep time (i.e., a third of 24-hour). Interestingly, for Group 2 we found three peaks of similar height around 8 hours prior to sleep, sleep onset, and sleep offset (Fig 3, C). We found the equivalent pattern when aligning by sleep offset (Fig 3, D). A similar pattern can be observed in the levels of plasma CORT over 24-hours, which displays a circadian rhythm that reaches a peak soon after awakening and a nadir during the night [37, 38].

### Mathematical modelling and the relationship between sleep, CORT and the distribution of EDs

To explore the hypothesis that sleep and CORT impact the distribution of EDs, we used a computational modelling framework. Within this framework, we assessed how changes to the overall excitability of brain regions impacted the overall likelihood of *in silico* EDs (see Materials and Methods for a detailed description of the model).

For Group 1, we used sleep staged polysomnography data collected from healthy controls (see Materials and Methods) as an external input to the excitability of the model *λ*_ext_. In Fig 4 Panel A we compared the model output for a virtual cohort of 66 individuals (in green) with the observed ED distributions for Group 1 (in blue).

**Fig 4.**
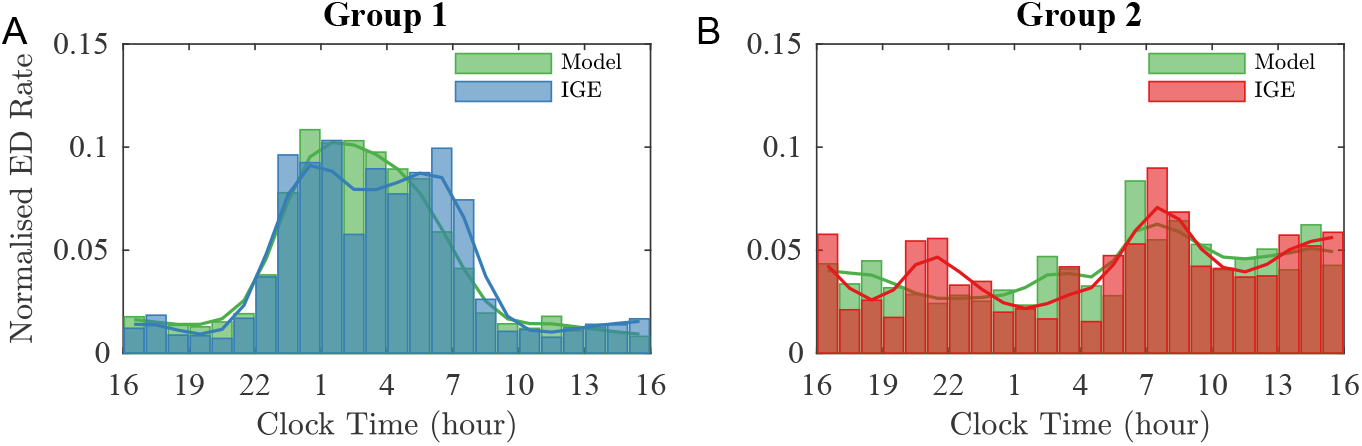
Model results compared with IGE data. (A) Histogram of EDs from Group 1 with IGE (blue) and histogram of EDs simulated using the model with *λ*_ext_ defined to mimic the different brain excitability during sleep stages (green). (B) Histogram of EDs from Group 2 with IGE (red) and histogram of EDs simulated using the model with *λ*_ext_ defined to mimic the impact of CORT on the brain excitability (green).

The model predicted a sharp increase in ED occurring during the first part of the sleep period, followed by a sharp decrease in the morning. The slow reduction in the number of EDs during the night is consistent with the observation that NREM sleep is predominant during the first part of the sleep, while REM is predominant during the second half [39]. Although the model captured most of the Group 1 ED variability (*R*^2^=0.9), it failed to capture the bimodal distribution in ED rate shown in the overnight data. It further failed to capture daytime variability in the ED rate, suggesting the presence of at least a second mechanism governing the ED propensity.

For Group 2, we used levels of CORT measured from healthy controls over the course of 24 hours (see Materials and Methods) as an external input to the excitability of the model *λ*_ext_. The model prediction for a virtual cohort of 41 individuals is shown in Fig 4 (in green). Comparing the model result with the data (in red), we found that the model captures the morning and afternoon peaks displayed by Group 2, although the latter occurs about an hour earlier in the model. We also note that the simulation does not account for the evening peak around 21:00. The overall variability explained by CORT in Group 2 is ∼60% (*R*^2^=0.59).

### Combined mechanism: sleep and CORT

For each group, we identified candidate mechanisms that could explain the majority of the observed distribution of EDs. However, we found that the model failed to capture some variability. For example in Group 1 the bimodal distribution during sleep, as well as some variability during the day, was not fully explained by the model. We therefore explored how combining the mechanisms of sleep and CORT impacted the ED distribution (see Materials and Methods). The strength of the influence of sleep and CORT is given by the parameters *p*_S_ and *p*_C_, respectively. Each parameter can vary from 0 (no impact on ED occurrence) to 1.5 (strong impact on ED occurrence). We used residual sum of squares (*RSS*) to identify the best fit (see Fig 5 and S4 Appendix for *R*^2^).

**Fig 5.**
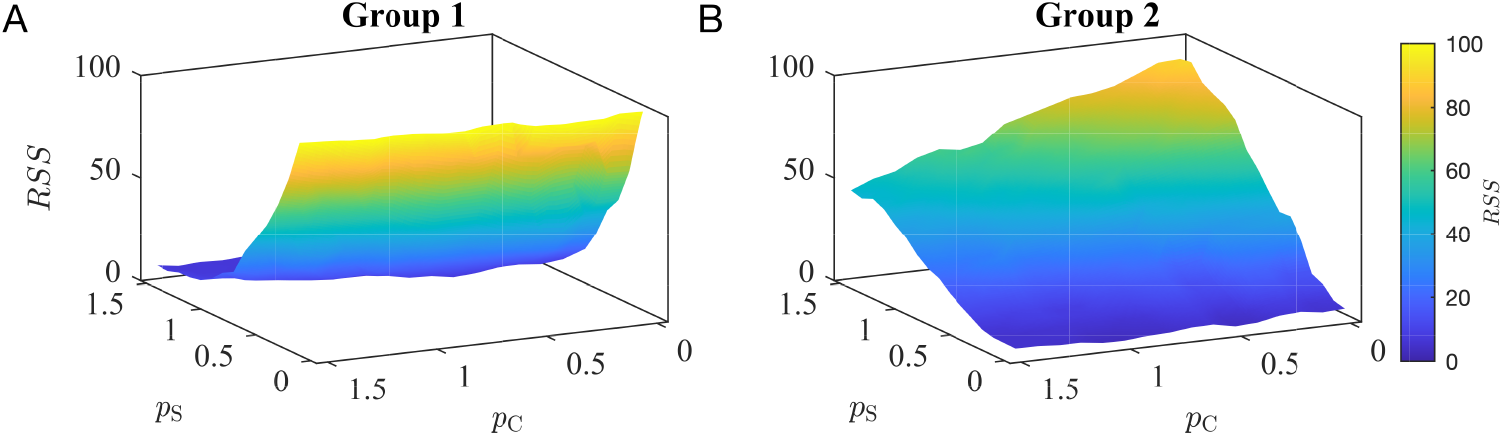
*RSS* values for the combined mechanism. Values of the residual sum of squares (*RSS*) computed over a grid of values of *p*_S_ and *p*_C_ for Group 1 (A) and Group 2 (B).

In Group 1, we found the best fit (lowest *RSS* values) was obtained when *p*_S_, *p*_C_ *>* 0 (Panel A). This result is consistent with our previous observation that sleep can explain the overnight peaks in EDs, with the contribution of CORT explaining variability during the day. This result suggests the coexistence of the two mechanisms (sleep and CORT) in Group 1. Fig 6 shows the model output corresponding to the lowest *RSS* for this group, which is when both sleep and CORT terms are present with *p*_S_ = 1 and *p*_C_ = 1.2. Combining these mechanisms increases the explained variability from 90% (only sleep) to 95% (*R*^2^ = 0.95).

**Fig 6.**
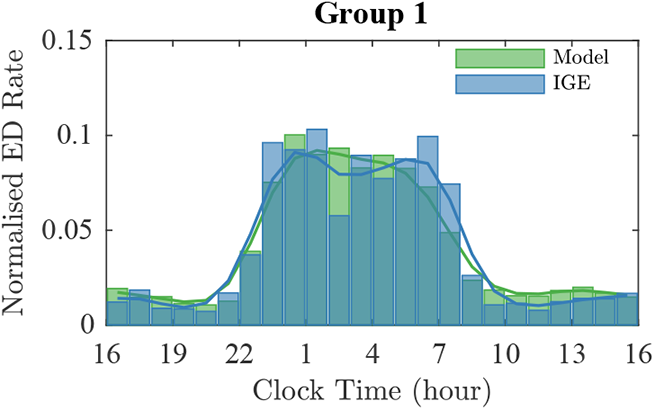
Best model fit for Group 1 compared with IGE data. Histogram of EDs from Group 1 with IGE (blue) and histogram of EDs simulated using the model with *λ*_ext_ defined to mimic the impact of the combined mechanism (sleep and CORT) on excitability (green). In this simulation, *p*_S_ = 1 and *p*_C_ = 1.2.

Conversely, the lowest values of *RSS* in Group 2 were obtained when *p*_S_ = 0 (Fig 5), suggesting that the best fit is obtained when CORT is the sole mechanism considered in the model (as in Fig 4, Panel B). Although additional mechanisms could be considered in explaining the remaining variance using this computational framework it is important to recognize the relatively modest sample-size of the remaining subgroups combined with the possibility of true random events (see S5 Appendix).

## Discussion

In this study, we provide a computational framework for assessing how the likelihood of epileptiform discharges is impacted by different physiological mechanisms and processes, such as sleep and stress. First, a data-driven analysis of the distributions of epileptiform activity from a large cohort of people with generalized epilepsies revealed the presence of two distinct groups within this cohort. To explain the underlying differences between these groups, we used a phenomenological mathematical model for simulating the activity of brain networks and excitability. Using this framework, we found that the patterns in the first group (Group 1) are strongly correlated with sleep, whereas the daily changes in ED likelihood in the second group (Group 2) can partially be explained by CORT. This framework provides an intuitive way of assessing the impact of external factors (e.g. sleep, stress, medication) on the overall likelihood of epileptiform activity, and can be used in the context of future experimental studies.

A data-driven approach was applied to the histograms of epileptiform discharges derived from 107 subjects with generalized epilepsies. First, we found that correlation and cluster analysis suggested the presence of two distinct groups within the overall cohort (of size 66 and 41 respectively). These two groups were not aligned with the clinical sub-types of IGE. Determining the periods of maximum ED likelihood in the two groups suggested sleep stages and CORT levels as candidate drivers for these ED distributions. EDs are increasingly understood as emerging from brain networks, with alterations to both the connectivity between brain regions, as well as the dynamics within regions, contributing to this emergence [40, 41]. In this regard, both sleep stage and levels of CORT have been shown to impact both functional connectivity [42, 43] and cortical excitability [44, 45]. Several studies have shown the correlation between sleep and epileptiform discharges [30, 32, 39] and how vigilance states may influence the likelihood of EDs in subjects with IGE [44, 46]. CORT is the main stress hormone in humans and its production and secretion are controlled by the hypothalamic-pituitary-adrenal (HPA) axis, the primary stress response system [18]. In stressful situations, the activity of the HPA axis increases, resulting in a higher secretion of CORT. In unstressed, basal conditions, cycling levels of CORT rise and fall over the day, with characteristic ultradian pulses [47]. This finding is consistent with the literature and self-reported data showing ED frequency increasing during the night time, early in the morning, and in stressful situations.

To investigate the impact of sleep and CORT on the ED likelihood during the day, we employed a phenomenological mathematical model to simulate brain excitability when perturbed by those external forces. Unlike in previous works where the variation of the brain excitability was constant [48] or perturbed by a fixed constant [49], this model describes cortical excitability as a dynamical variable that is modulated by dynamic external factors, such as sleep or CORT. We used sleep stages and CORT levels collected from healthy subjects to inform the dynamics of the variable representing the status of brain activity. However, in future work, the analysis should include CORT levels and sleep stage data derived from the EEG from the same individual, given that both of these variables show considerable inter-individual variability. Despite this limitation, our work shows a good fit between our model simulations and the observations. Indeed, we find that sleep accounts for 90% of the variability in Group 1 (*R*^2^ = 0.9) and CORT for ∼60% (*R*^2^ = 0.59) in Group 2.

Importantly, sleep alone cannot account for the changes in ED likelihood during wakefulness observed in Group 1. Furthermore, the model predicts a reduction in ED likelihood during the sleep time after an initial sharp increase during the first hours. This effect can be explained by the fact that NREM sleep, which is positively correlated to an increase of EDs, is predominant during the first third of the sleep period. However, the data shows an increase in ED occurrence before waking, which the model simulations fail to capture. Given that the level of CORT is known to increase around waking, this result suggests a combined effect of sleep and CORT. This result is quantitatively highlighted by the improvement in the accuracy of our model prediction when a combination of sleep and CORT have been considered and by the high percentage of variability explained by the combined model (95%, *R*^2^ = 0.95). It is important to emphasise that we only considered linear combinations, and future work could investigate a richer class of non-linear interactions and effects, especially given that sleep and stress themselves interact. This interaction may potentially lead to non-linear impacts on the likelihood of EDs.

Our model predicts peaks occurring during the day, for example one around 13:00 and one around 19:00, in Group 2. Those two peaks seem to occur a couple of hours earlier in the model than in the IGE cohort. The reason for such behaviour requires further investigation. One explanation could be additional physiological or behavioural drivers that we have not yet accounted for. Alternatively, it is important to highlight that CORT levels were measured in an independent control cohort. A future study would critically include simultaneous recordings of EEG and CORT, as well as detailed summaries of any anti-seizure treatment (e.g. timing and dose).

In summary, we provide a mathematical model as a tool to examine the role of external factors on the modulation of ED likelihood. We provide quantitative evidence that underlying physiological modulators for ED events exist. We identified sleep and CORT as such modulators by comparing our model predictions with data on ED events collected from IGE patients. Our choice of such factors is guided by the ED distribution in the EEG data and by previous studies investigating sleep and CORT, and the influence they have over the cortical excitability dynamics. Using only these two processes, we are able to account for the majority of the variability in the two groups. However, our results do not exclude other potential mechanisms affecting cortical excitability during the day, such as sleep deprivation or anti-seizure medication [27, 29]. Furthermore, other factors showing circadian rhythms, such as melatonin production or glucose levels, have also been shown to impact seizure incidence [50, 51, 51, 52]. Further research is needed to fully understand the overall mechanism underlying the modulation of ED events. In particular, simultaneous recordings of EEG and those factors are necessary to overcome the high intra- and inter-individual variability of the latter. Moreover, measurements should be taken from the same individual over prolonged periods, which would then inform the model framework (in particular the network structures and the excitability dynamics).

Ultimately, the modelling approach presented in this paper provides a framework to better understand what drives the occurrence of epilepsy-related activity observed in recordings of the brain.

## Materials and methods

### EEG data: Epileptiform Discharges in people with Idiopathic generalized Epilepsy

EDs were identified by an experienced EEG reader (U. S.) within EEGs from 107 people diagnosed with idiopathic generalized epilepsy (IGE). Scalp EEG recordings were collected for 24 hours using a 32-channel ambulatory EEG system (Compumedics Ltd.; Melbourne, Australia). Gold cup electrodes were attached with electrode paste according to the international 10-20 system. Subjects were encouraged to have at least seven to eight hours of night-time sleep prior to the EEG recording to guarantee optimum capture of ED. The study was approved by the Human Research Ethics Committees of St. Vincent’s Hospital and Monash Health. See [53] for more details.

### EEG data: Sleep-stages from healthy participants

Sleep-stages from 77 healthy participants were identified from EEG data collected at Monash University (Melbourne, Australia). Sleep polysomnography (PSG) was recorded across two consecutive nights in the laboratory (Compumedics Grael, Melbourne, Australia), using a bi-lateral 18-channel EEG, and two electro-oculographic (EOG, left and right outer canthi) and three electro-myographic (EMG, sub-mentalis) channels. EEG data were sampled at 512Hz. Sleep data for night 2 (following adaptation to the laboratory on night 1), were scored by a trained scorer, and in accordance with AASM criteria [54]. Monash University Human Research Ethics Committee approval was obtained for sleep study data (CF14/2790-2014001546; 2017-4204-11012; 2017-6008-8120; and 2020-5453-43401). We restrict our analysis to the 42 participants with sleep efficiency equal to or higher than 85% for night 2 (Table 1), as values less than this can be indicative of sleep disturbance.

**Table 1.** Characteristics of the subjects from the sleep cohort used in the simulations.

| Number | Female | Male | Mean Age (min, max) |
| --- | --- | --- | --- |
| 42 | 8 | 34 | 30.71 (18,64) |

### Blood data: CORT levels in healthy participants

Cortisol data was kindly provided by Elizabeth A. Young, University of Michigan. Blood samples for cortisol assay were collected from 6 healthy adult subjects via an intravenous catheter at 10 min intervals over a 24-hour period, as described previously [38, 55].

### Constructing a virtual cohort

A ‘virtual cohort’ approach was used to compensate for the differences in size and data modality across study groups (Group 1 (people with IGE): 66, Group 2 (people with IGE): 41, CORT (healthy participants): 6, sleep (healthy participants) : 42). In order to assess the potential impact of CORT and sleep on the distributions of EDs, new time series were sampled from the sleep and CORT data.

#### Sleep

To compensate for the smaller number of subjects in Group 1 compared to the sleep cohort, we randomly added 24 subjects (without repetition) from the sleep cohort. For Group 2, 41 subjects from the sleep cohort were randomly chosen from the 42 sleep participants.

#### CORT

To address the significant difference in group sizes (6 healthy participants vs 66 or 41 people with IGE), at each time-point (*i* = 1, …, 145), we used the synthetic minority oversampling technique (SMOTE) [56] to perform data augmentation. We therefore generated 60 synthetic CORT profiles for Group 1 and 35 synthetic CORT profiles for Group 2. SMOTE oversampling was performed with *k* = 3 (50% of the total) for the *k*-nearest neighbours used in the algorithm.

### Mathematical model

The model used in this study is based on the normal form of a subcritical Hopf bifurcation [48, 49, 57], whose co-existing states reflect two distinct types of neural activity. The first is a background state, represented by a steady-state solution in the model, whilst the second is an epileptiform state, represented by a high-amplitude oscillation. Transitions between these states are typically governed by either a white noise process or external perturbations. The model equations are given by:

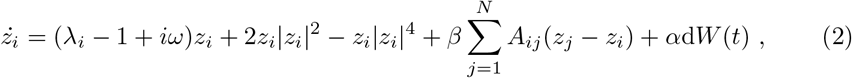

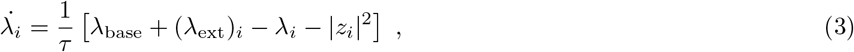

where *z*_*i*_(*t*) is dynamics of the *i*^*th*^ node (with *i* = 1, …, *N*), *W* (*t*) is a complex Wiener process, *λ*(*t*) is the excitability of node *i, λ*_base_ the baseline level of excitability, and *λ*_ext_(*t*) the external perturbations to the excitability. Typical parameter values for the model are given in Table 2, whilst *A* is an adjacency matrix, i.e. *A*_*i,j*_ is 1 if there is a connection between the *i*^*th*^ and *j*^*th*^ regions and 0 otherwise. For simplicity, all simulations were performed with a directed and connected 4-node graph (*N* = 4) (Fig 9), in line with [41].

**Table 2.** Parameters for the mathematical model.

| Parameter |  |  |
| --- | --- | --- |
| $\omega = 20 \text{ rad/s}$ | $\beta = 0.35$ | $\alpha = 0.055$ |
| $\tau = 3 \text{ s}$ | $\lambda_{\text{base}} = 0.65$ | |

**Fig 7.**
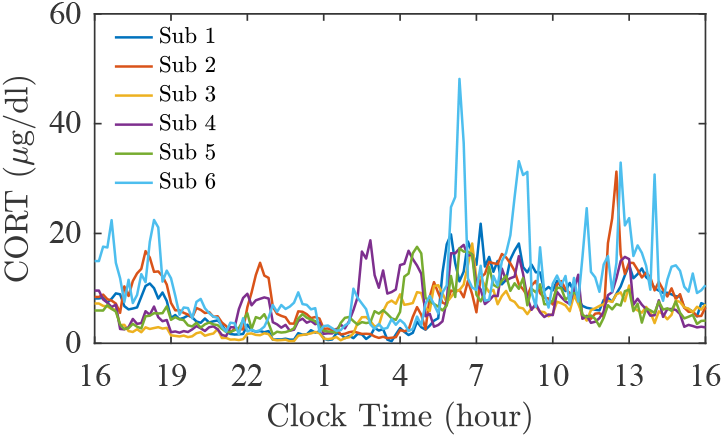
CORT 24-hour recordings. Blood samples for cortisol assay were collected from 6 healthy adult subjects via an intravenous catheter at 10 min intervals over a 24-hour period.

**Fig 8.**
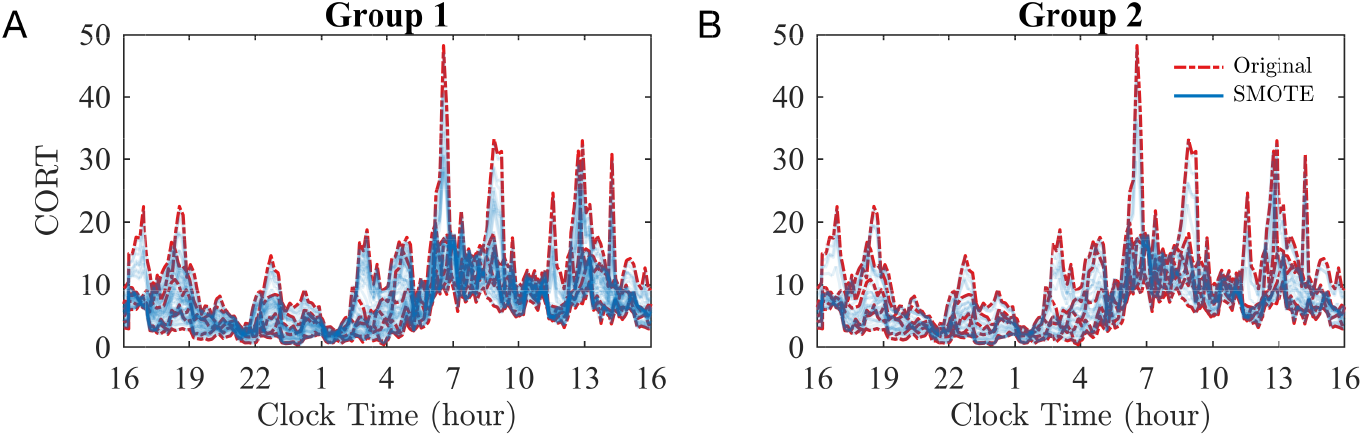
Synthetic CORT surrogates created using SMOTE. Original (red) and synthetic (blue) CORT profiles for Group 1 (A) and Group 2 (B). The synthetic data are obtained with the SMOTE oversampling algorithm with *k* = 3.

**Fig 9.**
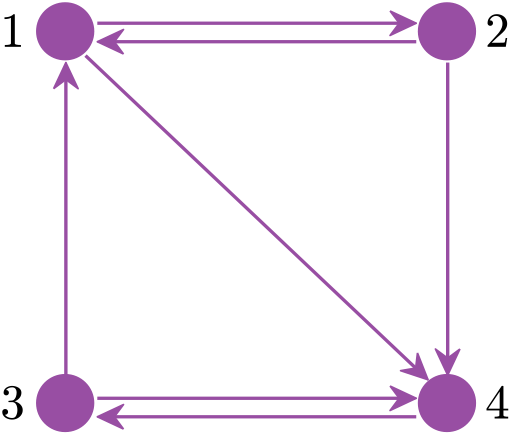
Schematic of the network used in the simulations. The network employed in the simulations is a directed and connected graph.

#### Influence of sleep

ED frequency has been observed to vary during sleep and to be higher during non-rapid eye movement (NREM), especially during stages N2 and N3, sleep which is associated with maximal synchronization, than during rapid eye movement (REM) sleep [58].

Therefore, we set *λ*_ext,sleep_ to its maximum value during the NREM state and to 0 during the REM phase. More precisely, *λ*_ext_ = 1 during N2 and N3, *λ*_ext_ = 0.5 during N1 and *λ*_ext_ = 0 during REM and wakefulness.

Panel A in Fig 10 illustrates a hypnogram representing the sleep stages recorded from a representative control participant (purple, top) and the corresponding rescaled *λ*_ext,sleep_ (black, bottom). The rescaling factor was found by optimising the simulated ED rate, when sleep is considered as external factor, and the data from Group 1. By minimising *RSS* we found the rescaling factor for sleep data to be *r*_S_ = 0.11.

**Fig 10.**
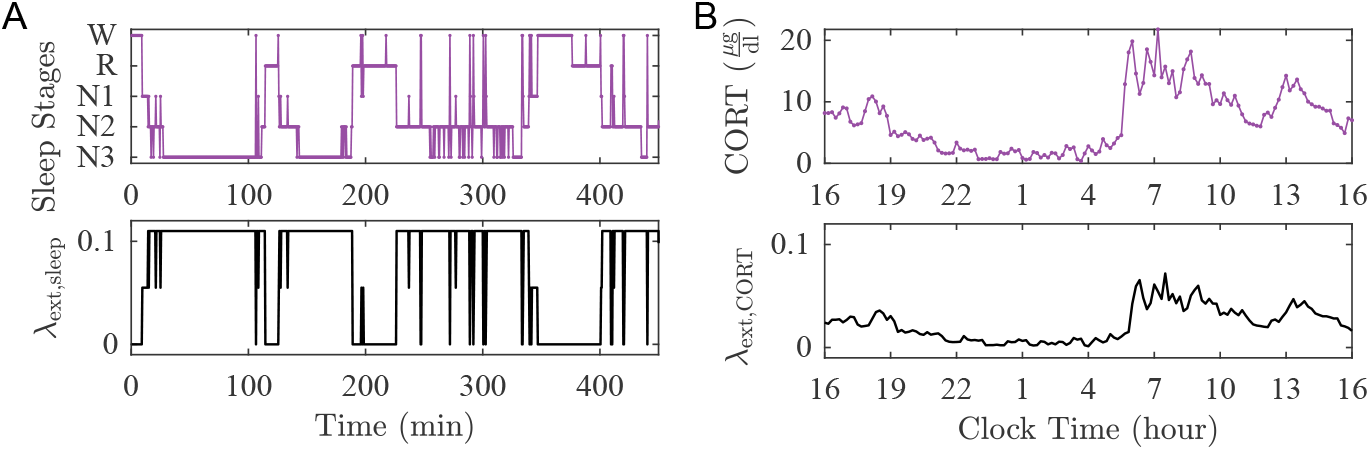
Modelling external perturbations informed by data. The external perturbation to brain excitability due to sleep, *λ*_ext,sleep_, and CORT, *λ*_ext,CORT_, were informed by using sleep stages (A) and CORT levels (B), respectively.

#### Influence of CORT

We defined *λ*_ext,CORT_ by rescaling the concentration values of CORT. Also, we account for the delay due to the non-genetic effect of CORT [59], by introducing a delay *τ* (min) in the *λ*_ext,CORT_ compared to the corresponding CORT profile. In our simulations, *τ* is from a normal distribution N (13, 5) [60, 61]. Panel B in Fig 10 illustrates the CORT profiles (purple, top) and the corresponding *λ*_ext,CORT_ (black, bottom). The rescaling factor was found by optimising the simulated ED rate, when CORT is considered as external factor, and the data from Group 1. By minimising *RSS* we found the rescaling factor for CORT data to be *r*_C_ = 0.0033.

#### Combined influence of sleep and CORT

To consider the combined effect of sleep and CORT, we defined

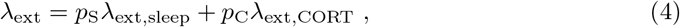

where *λ*_ext,sleep_ and *λ*_ext,CORT_ reflect the hypothesised physiological changes in brain excitability due to sleep and CORT, respectively.

Simulations were carried out over a grid where 0 ≤ *p*_S_ ≤ 1.5 and 0 ≤ *p*_C_ ≤ 1.5. For each parameter combination, we computed the residual sum of squares (*RSS*) to measure the discrepancy between the data and model predictions. More precisely, 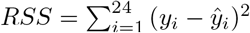, where *y*_*i*_ is the reported ED rate in the *i*^*th*^ 1-hour time interval, while *ŷ* _*i*_ is the model prediction for the corresponding time window.

## Acknowledgements

The authors acknowledge Dr. Charmine Diep for sleep staging of the clinical data, and members of the Monash Sleep and Circadian Sleep Laboratory for assistance in running the over-night studies.

## Author Contributions

I.M., W.W., J.R.T. conceived the study. I.M. performed formal analysis. U.S., C.A., S.L.L. contributed data used in the study. U.S., W.D., C.A., S.L.L., J.J.W., A.P.B., M.J.C. contributed to interpretation of clinical and experimental data. I.M., W.W. wrote the original draft. All authors contributed to the editing and reviewing of the manuscript.

## Supporting information

**S1 Appendix. Clustering and robustness of number of clusters**. The values of the Calinski-Harabasz criterion values for each number of potential clusters was tested for different numbers of bins.

**S2 Appendix. Impact of histogram bin width**. The size of the two clusters across the different bin width values is found to be consistent.

**S3 Appendix. Groups vs type of epilepsy**. Individuals with IGE are classified into childhood absence epilepsy (CAE), juvenile absence epilepsy (JAE), juvenile myoclonic epilepsy (JME), generalized epilepsy with generalized tonic-clonic seizures only (GTCSO), and genetic generalized epilepsy unspecified (GGEU), and divided into the two groups.

**S4 Appendix. *R***^**2**^ **statistic for the combined mechanism**. Values of *R*^2^ statistic computer over a grid of values of *p*_S_ and *p*_C_.

**S5 Appendix. Subgroups**. In both Group 1 and 2, identified two smaller sub-groups based on the similarity within their ED distributions.

